# Environmental monitoring for pathogen detection in zebrafish housing systems using molecular techniques: A 3Rs principles-based approach

**DOI:** 10.64898/2026.07.29.741654

**Authors:** Martina Angela Checco, Andrea Cacciamali, Silvia Dotti, Riccardo Villa

**Author notes:** Corresponding author: Martina Angela Checco.

## Abstract

Health monitoring is essential to ensure laboratory animal welfare and the reliability of experimental data in zebrafish research facilities. Conventional surveillance strategies based on resident or sentinel fish have limitations in terms of sensitivity and animal use, highlighting the need for alternative approaches consistent with the 3Rs principles. In this perspective, the present study evaluated the use of sludge collected from recirculating aquaculture systems as an environmental matrix for molecular health monitoring. Because sludge accumulates microorganisms and organic material from the entire system, it represents a promising sample for pathogen detection. Following an initial environmental surveillance phase to detect pathogens present in the system, the study aimed to optimise a molecular protocol for sludge analysis. To this end, four commercial DNA extraction kits were evaluated to assess their effectiveness in recovering bacterial DNA from sludge. Their performance was analysed by quantitative PCR in terms of extraction efficiency, repeatability, and limit of detection. The results highlighted the strengths and limitations of each DNA extraction protocol and confirmed the suitability of sludge as a non-lethal matrix for pathogen detection. These findings support the implementation of environmental monitoring as a practical and cost-effective alternative to sentinel-based surveillance, improving pathogen detection while reducing animal use in accordance with the 3Rs principles and Directive 2010/63/EU.

## 1. Introduction

Safeguarding animal welfare, along with the reliability and reproducibility of experimental data, plays a fundamental role in scientific research. In this context, health monitoring of laboratory animals represents an essential strategy, serving both as a Refinement and Reduction methodology within the framework of the 3Rs principles [1] and as a tool to improve the quality of experimental data [2]. Indeed, as with all biological systems, applying standardised procedures is the key to obtaining valid, reproducible, and comparable results.

The zebrafish (*Danio rerio*) is a vertebrate with biological, genetic, and experimental characteristics that make it an extremely powerful tool for biomedical research, particularly for *in vivo* studies, functional genetics, drug screening, and as a model for human diseases [3]. The growing interest in this species derives from its small size, high fertility, external and transparent embryonic development, and significant genetic homology with humans [4,5]. Furthermore, the development of methods that require increasingly limited employment of animals is leading to the use of models, such as fish, with a lower evolutionary level than mammals. This approach allows for the application of partial replacement, while simultaneously promoting scientifically valid and innovative research.

As occurs with other species used in animal testing, zebrafish colonies can also be susceptible to the emergence and spread of pathogens [6]. The presence of infections, particularly subclinical ones, represents a critical issue both for animal welfare and experimental research. Indeed, these infections can remain undetected for long periods, promote the spread of pathogens within the colony, lead to a progressive increase in mortality, and alter numerous biological parameters, including physical, immunological, and behavioural traits [6–8]

Furthermore, undetected infections can increase experimental variability and consequently require the use of a greater number of animals to achieve statistical significance, thereby raising significant ethical concerns. Health monitoring programs aim to monitor the presence of infectious and non-infectious diseases in the colony, inform and protect scientific research, facilitate the safe exchange of fish, and protect personnel (zoonoses). These objectives can only be achieved by implementing reliable health screenings and rigorous biosecurity protocols, as well as sharing health data among collaborating groups [8]

The Federation of European Laboratory Animal Science Associations (FELASA) periodically publishes updated guidelines for the microbiological monitoring of laboratory animals. These recommendations outline the fundamental aspects of health monitoring, specifying the pathogens to be screened, the biological samples to be analysed, the sampling frequency, and the most appropriate diagnostic techniques [2].

Microbiological monitoring of aquariums is based primarily on two approaches: sampling resident fish and sentinel fish. Resident fish that are euthanised for health reasons or found dead can provide useful information on the presence of pathogens in the colony, but their ability to detect low-prevalence infections is limited by the sample’s lack of representativeness and the genetic and experimental heterogeneity of the animals [6,8].

In recirculating systems, sentinel fish exposed to effluent water before filtration increase the likelihood of detecting pathogens shed by the colony and, if kept outside the circuit, prevent the reintroduction of infections. However, cumulative exposure can complicate the interpretation of results, while the use of sentinels exposed to water after filtration is generally uninformative and offers no advantages over direct sampling of fish from the colony [8].

Environmental monitoring represents a valid alternative to overcome these limitations. In addition to serving as a potential 3Rs strategy, it also offers important methodological advantages. Indeed, when combined with molecular techniques such as Polymerase Chain Reaction (PCR), environmental monitoring enhances pathogen detection sensitivity, reduces screening times, and lowers costs associated with animal management [9]

It is important to emphasise that the efficacy of environmental monitoring depends on the biological characteristics of the microorganism, its life cycle and its ability to persist or accumulate within the environment.

Several infectious agents affecting zebrafish may potentially be monitored through environmental matrices. Some parasites, such as *Pleistophora hyphessobryconis*, *Pseudoloma neurophilia*, and *Pseudocapillaria tomentosa*, are obligate pathogens, meaning they can reproduce only within a suitable host. Once their life cycle is complete, they are released into the environment as highly resistant spores or eggs. For obligate parasites, the level of environmental contamination depends directly on the number of spores or eggs released by infected animals [10].

For *Pseudoloma neurophilia*, several studies have shown that environmental DNA (eDNA) can be used to detect the microsporidian, although with varying sensitivity: the pathogen is released into the water intermittently, and continuous-flow systems significantly reduce the likelihood of detection [11,12].

For *Pleistophora hyphessobryconis*, on the other hand, there are currently no validated methods for eDNA detection. Diagnosis is based primarily on the identification of spores in tissues through histological examination or fresh specimens, while PCR on tissue samples is used as a confirmatory method. Therefore, monitoring of this microsporidian remains essentially based on the analysis of infected tissues [13,14]

For *Pseudocapillaria tomentosa*, tank detritus is an effective environmental sample, showing higher sensitivity than water or isolated faeces, as the nematode tends to accumulate in the organic matter deposited on the bottom. This makes detritus one of the few non-lethal samples with real diagnostic utility for this pathogen [10].

Among the infectious agents affecting zebrafish, mycobacteriosis is the bacterial disease most frequently encountered in experimental facilities [15].

Mycobacteria are ubiquitous opportunistic organisms, widely distributed in aquatic environments and capable of surviving both inside the fish and within biofilms present in rearing systems, which can serve as an important reservoir of infection [16,17]. Their ability to proliferate outside the host makes monitoring based exclusively on the analysis of animals or eggs insufficient for an accurate assessment of the colony’s health status [17]

In zebrafish, at least six species of *Mycobacterium* have been described: *M. abscessus*, *M. chelonae*, and *M. fortuitum* cause reduced survival and reproductive capacity [18], while *M. peregrinum*, *M. haemophilum*, and *M. marinum* are associated with more severe clinical forms and moderate to high mortality [19,20]. Several studies have shown that the detection of Mycobacterium spp. in environmental samples is more sensitive than that performed directly on zebrafish. In particular, the analysis of environmental samples, such as water, tank detritus, filtration system components, biofilms, and environmental swabs, has proven to be an effective approach to supplement and, potentially, replace monitoring based on sentinel fish [10,21,22]. Although environmental surveillance of zebrafish viral pathogens is still an emerging field, molecular detection of pathogens in these environmental matrices is technically feasible. However, relatively few studies have evaluated its application to zebrafish viruses, and its diagnostic performance has not yet been fully established. Consequently, environmental surveillance should currently be regarded as a complementary rather than a stand-alone approach for viral health monitoring [8].

Zebrafish used in biomedical research are reared in recirculating aquaculture systems (RAS), where water from the tanks undergoes filtration and disinfection processes before being reintroduced into the circuit. During the filtration phase, the so-called “sludge” accumulates—a matrix consisting of microorganisms, organic matter, and debris originating from the entire rearing system. This matrix was selected for the present study because the high concentration of biological material increases the probability of detecting any pathogens present in the system.

The present study aims to expand current knowledge on environmental monitoring as a potential alternative to monitoring based on the use of sentinel animals in zebrafish facilities. Specifically, this study investigates the use of environmental matrices for the detection of pathogens, including *Mycobacterium* species and *Pseudocapillaria tomentosa*, and simultaneously proposes molecular protocols for their identification. To achieve this goal, one of the major challenges is the development of a molecular approach that is sufficiently sensitive, specific, and robust for the analysis of complex environmental matrices. In particular, nucleic acid extraction and purification represent critical steps, as the quality and yield of the recovered genetic material directly influence the overall performance and reliability of PCR-based assays.

For this reason, four commercial DNA extraction kits, specifically designed for complex matrices and commonly used in pathogen research, were tested. These extraction methods were further compared through a sensitivity assessment aimed at identifying the most efficient protocol for the recovery of *M. marinum* DNA from sludge samples.

The results obtained encourage the use of an environmental monitoring strategy applicable to zebrafish facilities, capable of improving colony health surveillance, reducing the need for sentinel animals, and supporting the implementation of the 3Rs principles and Directive 2010/63/EU of the European Parliament and of the Council on the protection of animals used for scientific purposes [23].

## 2. Materials and Methods

### 2.1 Animal housing and sample collection

The housing rooms of the Istituto Zooprofilattico Sperimentale della Lombardia e dell’Emilia Romagna (IZSLER), where the zebrafish are reared, consist of two aquaria holding male and female fish, respectively. Within the aquaria, the water conductivity, pH, and temperature values are kept constant. Similarly, room temperature and levels of ammonia (NH_3_), nitrites (NO_2_), and nitrates (NO_3_) are strictly monitored and maintained. In the recirculating system, water from the tanks is collected and transported toward the treatment system. The first stage consists of a pre-filter, which retains larger solid particles such as uneaten feed, faeces, and suspended organic matter. After passing through the pre-filter, the water enters the main filter before passing through a UV lamp to reduce the microbial load. Following disinfection, the water is restored to optimal temperature and oxygenation conditions and is then directed back to the aquaria by the recirculation pump.

The aquarium pre-filter was provided by the facility operators during routine replacement, which occurred every fifteen days. Approximately 800 mg of sample were collected from the pre-filter, transferred into Eppendorf tubes, and refrigerated at +4 °C until the DNA extraction phase.

### 2.2 Environmental surveillance

This section describes the procedure used to detect pathogenic microorganisms in sludge samples. DNA was extracted using four different commercial kits, then amplified and sequenced to confirm the presence of *Mycobacterium spp*. Subsequently, species-specific PCRs were performed to identify *M. chelonae*, *M. marinum*, and the nematode *Pseudocapillaria tomentosa*.

#### 2.2.1 DNA extraction

Approximately 200–250 mg of collected sludge was used for DNA extraction. Pathogen DNA was extracted using the following commercial kits: Quick-DNA Fecal/Soil Microbe Miniprep (Zymo Research, Irvine, CA, USA), NucleoSpin® DNA Stool (Macherey-Nagel GmbH & Co. KG, Düren, Germany), ReliaPrep™ Blood gDNA Miniprep System (Promega Corporation, Madison, WI, USA), and DNeasy® PowerSoil® Pro Kit (Qiagen, Hilden, Germany). Detailed information for each kit is presented in Table 1.

**Table 1:** Characteristics of the nucleic acid extraction kits evaluated in this study.

| Extraction kit | Manufacturer | Size | Extraction principle | Sample size | Thermic treatment for lysing sample | Final DNA volume |
| --- | --- | --- | --- | --- | --- | --- |
| Quick-DNA<br>Fecal/Soil<br>Microbe<br>Miniprep | Zymo | 50 Preparations | Silica<br>membrane-<br>based column | ≤250 mg | None | 50-100 µL |
| ReliaPrep™<br>Blood gDNA<br>Miniprep<br>System | Promega | 50 Preparations | Bead-beating,<br>Silica<br>membrane-<br>based column | 250mg | 56°C-10 min | 100 µL |
| NucleoSpin®<br>DNA Stool | Machery Nagel | 50 Preparations | Silica<br>membrane-<br>based column | 180-220mg | 70°C-10min | 30–100 µL |
| Dneasy Power<br>Soil Pro | Qiagen | 50 Preparations | Silica<br>membrane-<br>based column | 100- 200 mg | None | 50-100 µL |

#### 2.2.2 DNA amplification

DNA extracted from the sludge samples was subjected to PCR analysis for the detection of *Mycobacterium spp*. The primers and probes used for targeting the 16S ribosomal RNA (rRNA) gene are reported in Table 2. A PCR master mix was prepared in a 24.5 µL reaction volume containing 2.5 µL of template, 12.2 µL of GoTaq® G2 Green Master Mix (Promega), and a 5 µM primer set (Metabion International AG, Planegg, Germany). The reaction was conducted using a T100™ Thermal Cycler (Bio-Rad Laboratories, Hercules, CA, USA) with a thermocycling profile consisting of an initial denaturation, followed by 40 cycles of 95°C for 30 s, 65.5°C for 30 s, and 72°C for 70 s.

**Table 2.**
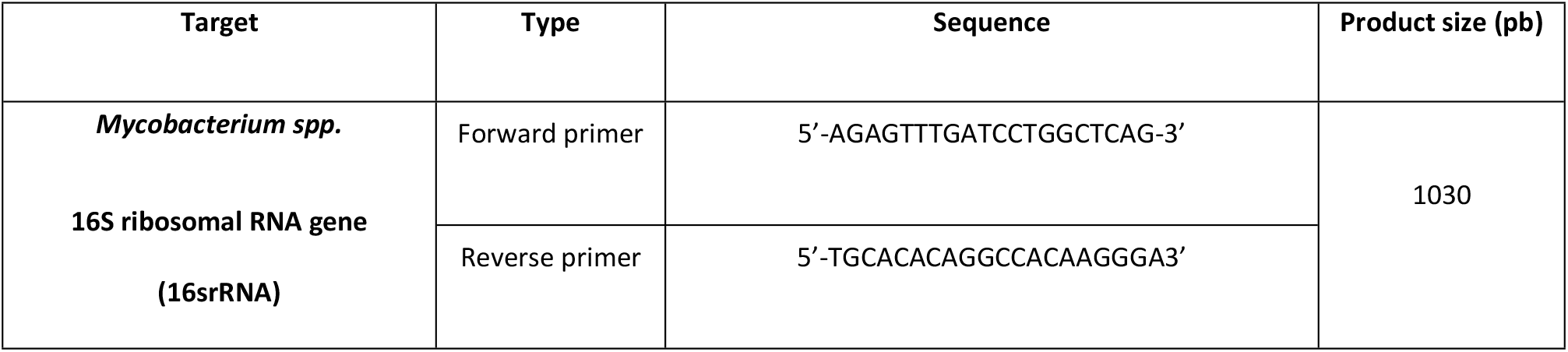
Primers used for the PCR assay for the detection of Mycobacterium spp.

To detect the amplified products, the QIAxcel® Connect Capillary Electrophoresis Instrument (Qiagen) was utilised. The procedure was performed in strict accordance with the manufacturer’s instructions.

#### 2.2.3 DNA Sequencing

PCR-positive amplicons were purified and subjected to Sanger sequencing to confirm the specificity of amplification. Briefly, to eliminate the amplification primers, the amplicon was treated with Exonuclease I (Exo I) (Thermo Fisher Scientific, Waltham, MA, USA). To remove unincorporated dNTPs from the amplification reaction, a dephosphorylation treatment was performed using Thermosensitive Alkaline Phosphatase (FastAP) (Thermo Fisher Scientific). The reaction mixture contained 0.5 µL of Exo I (20 U/µL) and 1 µL of FastAP (1 U/µL). The thermal cycler profile consisted of an incubation at 37°C for 15 minutes, followed by an enzymatic inactivation phase at 85°C for 15 minutes. To set up the sequencing reaction, the Forward and Reverse Sequence Mixtures and the BigDye™ Terminator v3.1 Cycle Sequencing Kit (Thermo Fisher Scientific) were used. For each mixture, the following reagents were added: 1.5 µL of Ready Reaction Mix, 3.2 pmol of Forward or Reverse Sequencing Primer, and 1 µL of 5X Sequencing Buffer. The reaction was performed under the following thermocycling conditions: 96°C for 90 s, followed by 25 cycles of 96°C for 10 s, 55°C for 5 s, and 60°C for 4 minutes. For product analysis, the sample was purified and transferred to the SeqStudio™ Genetic Analyser (Applied Biosystems, Thermo Fisher Scientific, Waltham, MA, USA) for capillary electrophoresis.

The resulting sequences were compared with reference sequences available in GenBank using BLAST to verify their assignment to the genus Mycobacterium spp.

#### 2.2.4 Species-specific qPCR

DNA extracted from the sludge was subjected to Real-Time PCR (qPCR) amplification for the detection of *M. chelonae*, *M. marinum*, and *Pseudocapillaria tomentosa*. The specific primers and probes used in this study are reported in Table 2 [24,25]. For the detection of the target genes, a qPCR master mix was prepared in a 20µL final reaction volume containing 3µL of template DNA, 5µL of GoTaq® Enviro qPCR Master Mix (Promega), 0.4µM of each primer (Metabion), 0.2µM of the species-specific probe (Metabion). The amplification was performed using a CFX96™ Real-Time PCR Detection System (Bio-Rad) under specific thermocycling conditions optimised for each pathogen. For *M. chelonae*, the thermocycling profile consisted of an initial denaturation at 95°C for 10 minutes, followed by 40 cycles of denaturation at 95°C for 15 seconds and annealing/extension at 61.5°C for 1 minute. For *M. marinum*, the thermocycling profile consisted of 95°C for 5 min, followed by 40 cycles of 95°C for 0.15 s and 60°C for 45 s. For *Pseudocapillaria tomentosa* detection, the thermocycling profile involved an initial denaturation at 95°C for 5 minutes, followed by 40 cycles of denaturation at 94°C for 40 seconds, annealing at 65°C for 30 seconds, and a final extension at 72°C for 1 minute and 30 seconds.

#### 2.2.5 Statistical Analysis

For each extraction kit, both DNA extraction and PCR amplification were performed in duplicate in two independent experiments. Results are presented as the mean ± standard deviation (SD).

### 2.3. Analytical validation

The following paragraphs describe the procedure used to evaluate the sensitivity of the DNA extraction kits employed in this study. Bacterial DNA was extracted from the sludge samples, and qPCR was performed to assess extraction efficiency, repeatability, and the limit of detection (LOD), defined as the lowest concentration at which 95% of samples yielded a positive result.

#### 2.3.1 Bacterial preparation for study sensitivity of DNA extraction kit

*M. marinum* was obtained from the Biobanking of Veterinary Resources (BVR) (IZSLER).

Bacterial cultures were prepared by the IZSLER Specialised Bacteriology Laboratory (Brescia, Italy). Frozen stocks were thawed and propagated in Middlebrook M7H9 medium. Tenfold serial dilutions were prepared and plated on Middlebrook M7H10 agar. Colonies were counted after incubation at 37°C for 20 days.

#### 2.3.2 Spiking of the environmental sample

Environmental matrices were collected from IZSLER experimental animal facilities (Brescia, Italy).

Approximately 200-250 mg of sludge collected from the aquarium pre-filter was inoculated with 20 µL of an *M. marinum* suspension prepared in phosphate-buffered saline (PBS) at concentrations ranging from 10² to 10⁷ CFU/mL.

#### 2.3.3 DNA extraction and amplification

To evaluate the DNA extraction performance for each kit, 25 µL of the Internal Control (IC) DNA (High Conc.) (Qiagen) was added to each dilution. In addition, a negative control (uninoculated matrices) and a positive control (matrices spiked with *M. marinum*) were included in every experimental run.

Pathogen DNA was extracted with the following diagnostic kits: Quick-DNA Fecal/Soil Microbe Miniprep (Zymo), NucleoSpin**®** DNA Stool (Macherey-Nagel™), Reliaprep™ Blood gDNA miniprep System (Promega), Dneasy Power Soil (Qiagen). Information about each kit is shown in Table 1.

For the detection of the bacterial gene, a qPCR master mix was prepared in a 20 µL reaction volume containing 3 µL of template, 5 µL GoTaq(R) Enviro qPCR System (Promega), primer set 0.4 µM (Metabion), species-specific probe 0.2 µM (Metabion), and 2 µL of IC assay Quanti Fast Pathogen (Qiagen). The reaction was conducted using a CFX96 Real-Time PCR System (Bio-Rad) with a thermo-cycling profile consisting of 95°C for 5 min, followed by 40 cycles of 95°C for 0.15 s and 60°C for 45 s.

Primers and probes used for *M. marinum* detection are based on the bacterial ERP gene, a species-specific region in the *Mycobacterium* genome [24].

The Ct values were used to assess reproducibility and the limit of detection (LOD). The LOD for each DNA extraction kit was defined as the lowest concentration of the spiked sample at which 95% of the samples yielded positive results. Positive and negative controls were included in the qPCR reaction plates.

#### 2.3.4 Statistical Analysis

For each extraction kit, the test was performed in triplicate, while the amplification was executed in six replicates. A statistical evaluation was performed to compare Ct values across different commercial kits using one-way analysis of variance (ANOVA), followed by Student’s t-test. Differences were considered statistically significant when the p-value (P) was ≤ 0.05. Prism-GraphPad was used to analyse and plot the replicate experiments, and the results are presented as mean ± standard deviation (SD). In addition, the coefficient of variation (CV%) of the Ct values was calculated to assess the repeatability of the measurements and the consistency of the extraction-PCR workflow.

## 3. Results

### 3.1 Environmental surveillance

### 3.1.1 Pathogen Detection

PCR analysis revealed contamination with *Mycobacterium spp*. Subsequent Sanger sequencing confirmed these results and produced electropherograms with mixed sequences, showing 97–98% identity with *Mycobacterium spp*. The qPCR confirmed the presence of *M. chelonae* and *Pseudocapillaria tomentosa* in the prefilter, but not of M. *marinum*. The Ct values for each detected pathogen obtained with the different DNA extraction kits are presented in Table 3.

**Table 3.**
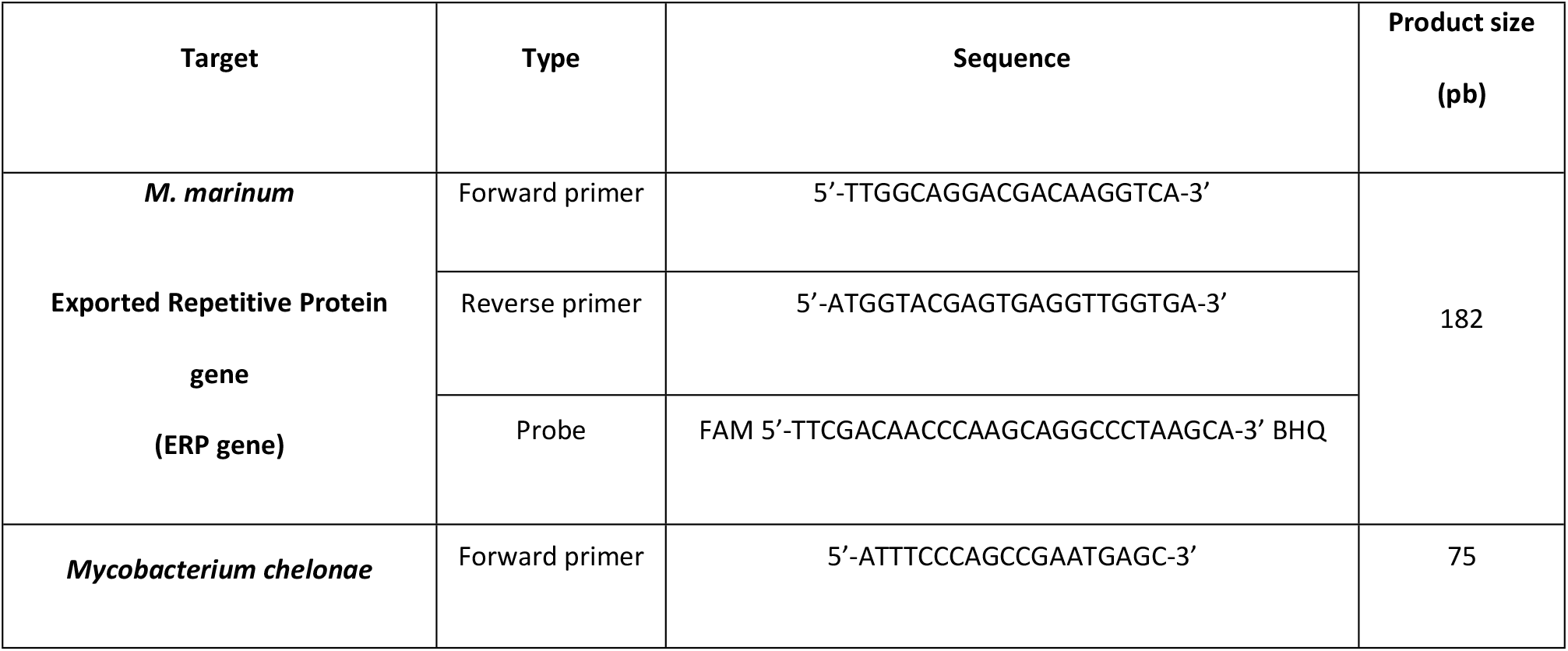
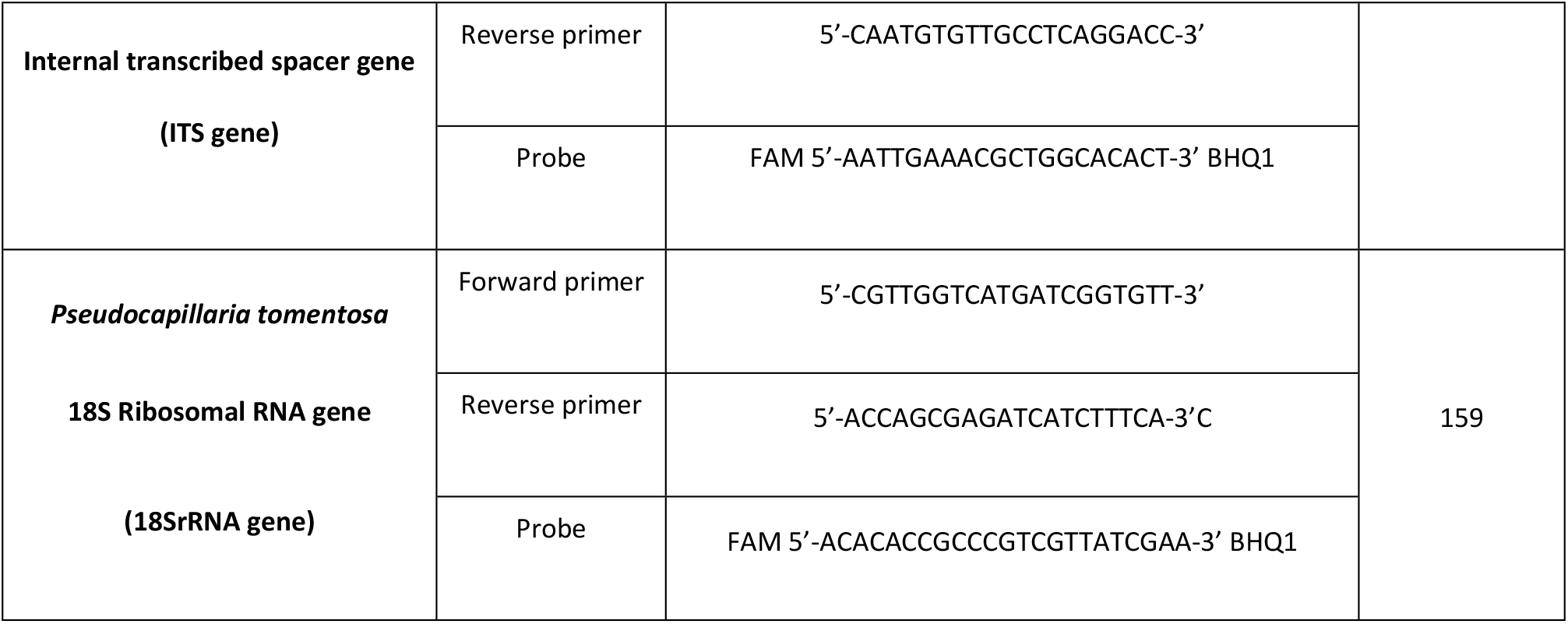
Primers used for the qPCR assays for the detection of target pathogens.

**Table 3.** Ct values for the detection of *M. chelonae*, *M. marinum* and *Pseudocapillaria tomentosa* in environmental samples collected from aquarium filters and prefilters. (N/A: not available)

| Microorganism surveillance | Extraction kit | Ct Avg (Mean ±SD) |
| --- | --- | --- |
| *M. chelonae* | Quick-DNA Fecal/Soil Microbe Miniprep Kit (Zymo) | 36.8±0.1 |
|  | Relia Prep Blood gDNA (Promega) |  |
|  | NucleoSpin™ DNA Stool (Macherey-Nagel™) | 32.09 ±2.8 |
|  | Dneasy Power Soi (Qiagen)I | 29.2±2.07 |
| *M. marinum* | Quick-DNA Fecal/Soil Microbe Miniprep Kit (Zymo) | N/A |
|  | Relia Prep Blood gDNA (Promega) | N/A |
|  | NucleoSpin™ DNA Stool (Macherey-Nagel™) | N/A |
|  | Dneasy Power Soi (Qiagen)I | N/A |
| *Pseudocapillaria tomentosa* | Quick-DNA Fecal/Soil Microbe Miniprep Kit (Zymo) | 35±0.7 |
|  | Relia Prep Blood gDNA (Promega) | N/A |
|  | NucleoSpin™ DNA Stool (Macherey-Nagel™) | 35.6 ±0.06 |
|  | Dneasy Power Soi (Qiagen)I | N/A |

### 3.2 Analytical validation

#### 3.2.1 DNA extraction efficiency

To compare the bacterial nucleic acid extraction efficiency of each kit, *M. marinum* was spiked into environmental samples. For *M. marinum* detection, bacteria were added to sludge obtained from aquarium pre-filters at concentrations ranging from 1×10^7^-to 1 x10^2^ CFU/mL. The DNA samples were then analysed by qPCR. To support efficient amplification of the target samples and controls, the sludge matrix was diluted 20-fold, and the IC volume was increased to 25 µL before extraction.

Following minor adjustments to the extraction procedure, successful amplification of the internal controls and the analysed target was achieved for all tested kits, demonstrating the recovery of amplifiable DNA from the environmental matrix.

#### 3.2.2 Comparison of the extraction kits by qPCR for *M. marinum* detection

Table 4 shows the qPCR limit of detection (LOD) based on DNA extracted from aquarium pre-filter sludge. The lowest LOD detected by the Quick-DNA Fecal/Soil Microbe Miniprep (Zymo) extraction kit was 10^2^ CFU/mL. The other LODs were detected for the NucleoSpin**®** DNA Stool (Macherey-Nagel™), Dneasy Power Soil (Qiagen) and ReliaPrep™ Blood gDNA Miniprep System (Promega) kits at 10^4^ CFU/mL, 10^6^ CFU/mL, and 10^7^ CFU/mL, respectively.

**Table 4.** Limit of detection values obtained following qPCR analysis for the identification of the target *M. marinum* from sludge samples taken from the pre-filter of the aquariums.

| Extraction kit | Sludge |  |
| --- | --- | --- |
| | LOD (CFU/mL) | Ct Avg (Mean ±SD) |
| Quick-DNA Fecal/Soil Microbe Miniprep (Zymo) | $1 \times 10^2$ | $32.10 \pm 2.1$ |
| ReliaPrep™ Blood gDNA Miniprep System (Promega) | $1 \times 10^7$ | $35.01 \pm 0.7$ |
| NucleoSpin® DNA Stool (Machery Nagel) | $1 \times 10^4$ | $34.87 \pm 2.4$ |
| Dneasy Power Soil Pro (Qiagen) | $1 \times 10^6$ | $30.26 \pm 2.3$ |

Figure 1A presents the distribution of Ct values from *M. marinum* DNA extracted at a concentration of 1×10⁷ CFU/mL using the different kits. The lowest and highest average Ct values were recorded for the Quick-DNA Fecal/Soil Microbe Miniprep (Zymo), Miniprep System (Promega) kits, with means of 26.75 and 35.21, respectively. The average Ct values obtained from the amplification of DNA extracted with the other kits tested were 29.69 for NucleoSpin**®** DNA Stool (Macherey-Nagel™), and 28.14 for Dneasy Power Soil (Qiagen), respectively. Differences in Ct values between kits used were significant by one-way (P≤0.05). In particular, pairwise comparison revealed a significant difference between Quick-DNA Fecal/Soil Microbe Miniprep (Zymo) and the other kits analysed (P≤0.05).

**Figure 1:**
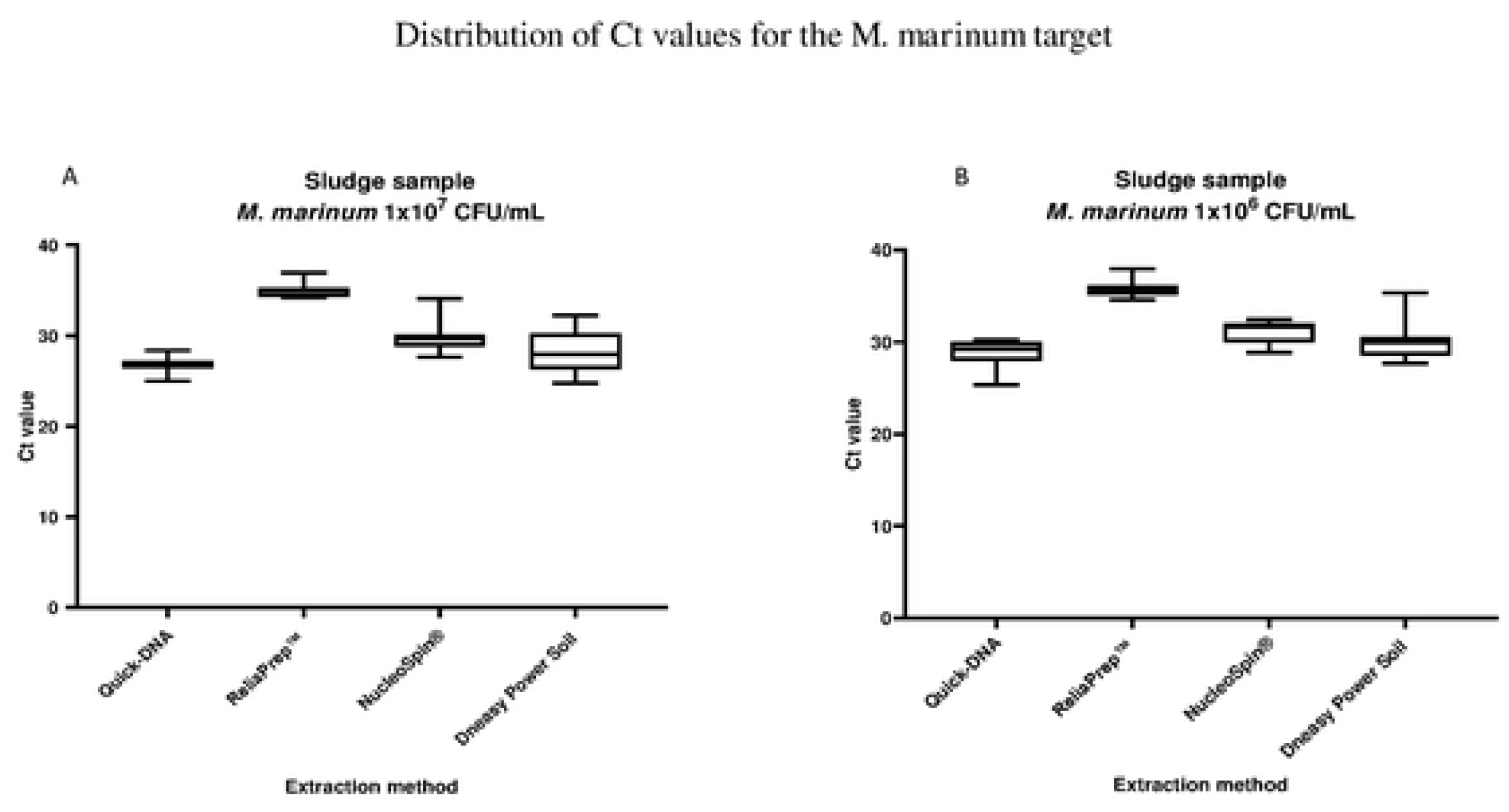
The graphs show the distribution of Ct values for the M. marinum target extracted with four different extraction kits from a sludge sample taken from the aquarium pre-filter. The comparison between the different extraction kits showed significant differences (p≤0.05, n=18) for both bacterial concentrations considered (1×10⁷ CFU/mL, 1×10⁶ CFU/mL).

In addition, Figure 1B shows the Ct value distribution at 1×10⁶ CFU/mL. The Quick-DNA Fecal/Soil Microbe Miniprep (Zymo) kit again yielded the lowest mean Ct value (28.8), while the Reliaprep™ Blood gDNA Miniprep System (Promega) had the highest (35.7).

ANOVA analysis confirmed statistical differences (P≤0.05), except between Dneasy PowerSoil (Qiagen) and NucleoSpin**®** DNA Stool (Macherey-Nagel™), which showed similarities.

The repeatability of results was assessed by calculating the coefficient of variation (CV%). Quick-DNA Fecal/Soil Microbe Miniprep (Zymo), NucleoSpin**®** DNA Stool (Macherey-Nagel™), and ReliaPrep™ Blood gDNA Miniprep System (Promega) kits demonstrated good repeatability (CV≤ 5%). The ReliaPrep™ Blood gDNA Miniprep System (Promega) kit, in particular, showed the lowest (CV∼2%) across tested concentrations.

## 4. Discussion and Conclusions

The development of monitoring plans that include environmental sampling and the use of molecular methods to protect the health of laboratory animals plays a central role in the application of the 3Rs principle [1]. Environmental monitoring contributes to the Refinement principle by enabling less invasive surveillance that better respects animal welfare, as well as to the Reduction principle by providing complementary information to that obtained from sentinel animals and allowing early detection of subclinical conditions that could increase experimental variability. In this context, analysing environmental matrices represents a promising approach to complement or, in some cases, replace traditional animal-based methods.

This project aimed to develop a new diagnostic approach for zebrafish to characterise the pathogens circulating within the facility environment. The environmental monitoring plan focused on three key assessment points:

1. The type of sample to be collected, to identify the most suitable matrix for the detection of different pathogens;
2. The presence of specific microorganisms within the matrix, assessed using targeted molecular methods;
3. The enhancement of molecular extraction and amplification performance, necessary to optimise diagnostic sensitivity in a complex matrix.

The entire study was therefore structured to progressively address each of these aspects, with the ultimate goal of defining a reliable and applicable diagnostic protocol for zebrafish systems.

The sample type was selected based on currently available knowledge regarding environmental resources in aquariums. Water represents a commonly used matrix for pathogen screening in aquatic systems [10,21,26]; however, our preliminary tests revealed that it is poorly suited for detecting pathogens present at low concentrations. Furthermore, the concentration step using specific filters—which is required to achieve adequate diagnostic sensitivity—involves very long processing times that are hardly compatible with routine monitoring requirements.

Pre-filter sludge represents a particularly interesting matrix for health monitoring in zebrafish facilities because it constitutes a composite sample of the entire recirculating aquaculture system. During normal system operation, the pre-filter retains faecal material, biofilm, organic debris, and microorganisms originating from all connected tanks, thereby concentrating over time any pathogens that may be present [10,27,28]. Consequently, sludge can offer a higher detection probability compared to sampling water or individual animals, especially when infection is present at a low prevalence or in a subclinical phase.

An additional advantage is the non-invasive nature of this sampling method. Sludge collection can be carried out during routine system maintenance without handling or sacrificing any fish, thereby contributing to the reduction of sentinel animal use and aligning with the 3Rs principles. Furthermore, because sludge is a matrix that accumulates over time, it provides an integrated overview of the system’s microbiological contamination rather than a single snapshot limited to the time of sampling.

During the environmental surveillance phase of the study, filter sludge samples were collected as part of routine aquarium maintenance, with aquarium filters being replaced every 15 days. Although a rigid experimental plan was not established for sample collection, this approach closely reflects the real-world operation of recirculating systems, while avoiding the DNA degradation issues observed in filters left in the sumps for extended periods [22]

It is important to emphasise that monitoring methods based on environmental samples have inherently variable sensitivity, which is influenced by both the pathogen type and the nature of the sample being analysed. Not all etiological agents can be detected with equal effectiveness, as they differ in the amount of genetic material and biological particles released into the environment, in the stability of their DNA (or RNA) once shed into the aquatic system, and in the pathogen’s distribution across the liquid phase, suspended solids, and biofilm. Furthermore, the life cycles and shedding patterns vary considerably among microorganisms, thereby affecting the overall sensitivity of environmental sampling [29]

When compared with the literature, our approach demonstrates clear consistency in the detection of mycobacteria in the environment [10,21,22,30,31] but a significant advantage in the detection of *Pseudocapillaria tomentosa.* From this perspective, our study and Crim MJ et al. 2017 can be considered methodologically similar, as both are based on environmental matrices rich in detritus and faecal material, which are the most suitable for detecting pathogens not suspended in the water column [10]. Subsequent studies have instead shown very low or no sensitivity in environmental samples, since filtered water or sump filters do not capture heavy faecal material [21,22].

The techniques used to detect environmentally-acquired infections play a crucial role in health surveillance programs. From a methodological standpoint, sludge presents certain challenges. It is a complex matrix, rich in substances that can potentially inhibit PCR, and it also contains high amounts of non-target environmental DNA. For this reason, the selection of nucleic acid extraction and amplification protocols is a crucial step in the development of a reliable diagnostic protocol.

The detection of pathogens belonging to the genus Mycobacterium spp. was performed by amplifying the 16S rRNA gene, revealing the presence of mycobacteria in samples extracted using various purification kits. Subsequent sequencing of the amplicon using the Sanger method provided more detailed information than conventional PCR alone. In particular, the chromatograms showed the presence of double peaks, suggesting the coexistence of multiple sequences in the same sample, while BLAST analysis yielded identity values ranging from 97% to 98% compared to reference sequences deposited in GenBank. These results are likely attributable to the heterogeneous nature of the sludge, which contains DNA from numerous microorganisms that may have been co-amplified during PCR. Furthermore, the high conservation of the 16S rRNA gene limits identification to the species level, and in environmental samples, the presence of mixed bacterial communities and the quality of the extracted DNA can reduce alignment quality and sequence identity compared to what is observed in pure isolates [32].

For this reason, the other pathogens under investigation were not sequenced; instead, their identification was performed using species-specific qPCR, which allowed for the reliable differentiation of *M. chelonae* and *M. marinum* and the detection of *Pseudocapillaria tomentosa*. Overall, these results highlight how the inherent complexity of environmental matrices makes it necessary to search for the correct molecular approach to achieve accurate pathogen identification.

The qPCR tests confirmed the presence of *M. chelonae* and *Pseudocapillaria tomentosa* in the sludge samples collected from the IZSLER aquaria. However, the recorded Ct values were generally high or, in some cases, undetectable, and varied depending on the extraction kit used. These findings are likely attributable to the low concentration of target DNA and/or the presence of PCR inhibitors that reduced amplification efficiency. Such results are consistent with the characteristics of complex environmental matrices and should therefore be interpreted in the context of the nature and complexity of the samples. It is well known, in fact, that high-yield extraction methods can co-purify PCR inhibitors [33], while diagnostic performance can vary depending on both the extraction kit used and the biological characteristics of the pathogen being detected [34]. Overall, the high Ct values observed highlight a methodological limitation that can be mitigated by optimising DNA extraction and purification procedures. The second part of the study focused on identifying the most suitable nucleic acid extraction protocol for this complex environmental matrix. To this end, the four extraction kits selected during the environmental surveillance phase were compared and evaluated in terms of efficiency, sensitivity, and reproducibility. To compare the recovery of bacterial nucleic acid, different concentrations of *M. marinum* were added to the environmental samples. Specifically, sludge collected from the aquariums’ pre-filters was inoculated with bacterial suspensions ranging from 10⁷ to 10² CFU/mL; the DNA was then extracted and subsequently amplified using species-specific qPCR.

The efficiency of the extraction procedure was inferred from the successful detection of the internal control and the analysed targets. Their amplification indicated that the extracted DNA was of sufficient quality for molecular analysis and that all the tested kits enabled the recovery of amplifiable DNA from the environmental matrix. Among the kits tested for the detection of *M. marinum*, the Quick DNA Fecal/Soil Microbe Miniprep (Zymo) proved to be the most sensitive, with a significantly lower LOD than that of the other kits. This result suggests that PCR inhibitors, which have a decisive influence on the test’s sensitivity, are removed more effectively using this extraction kit, thereby improving the ability to detect low concentrations of target DNA.

The reproducibility of the results was assessed by calculating the coefficient of variation (CV%), which showed differences depending on the kit used. The CV of Ct values serves as an indicator of the repeatability of the overall extraction-PCR workflow, reflecting the consistency of the DNA recovered by the extraction method as assessed indirectly through qPCR. Lower variability in Ct values suggests a more reproducible extraction process, assuming constant PCR conditions. The Quick DNA Fecal/Soil Microbe Miniprep (Zymo), NucleoSpin® DNA Stool (Macherey Nagel™), and Reliaprep™ Blood gDNA Miniprep System (Promega) kits demonstrated good reproducibility (CV ≤ 5%). In particular, the Reliaprep™ Blood gDNA Miniprep System (Promega) exhibited the lowest CV (approximately 2%) across the various concentrations tested, indicating remarkable operational stability despite the limitations observed in terms of diagnostic sensitivity.

In conclusion, the results of this study can assist laboratories in selecting the most appropriate in selecting the most appropriate molecular protocol according to appropriate molecular protocol according to sample type, matrix complexity, and the nature and abundance of the target pathogens. Identifying the optimal matrix, along with refining extraction and amplification protocols, can significantly improve the use of environmental samples for diagnosing pathogens in aquaria facilities, thereby reducing the need to sacrifice sentinel animals. This approach is fully in line with international guidelines and the new paradigm aimed at protecting laboratory animals.

## Acknowledgments

We are grateful to Dr.ssa Marta Consoli for the excellent work done in this project and to IZSLER Experimental Animal Facility and IZSLER Specialised Bacteriology Laboratory for the materials provided.

## Author Contribution Statement

Each author provides their contribution.

In the following lines, contributions are reported

Martina Angela Checco: Conceptualisation; Data curation; Investigation; Methodology; Writing - original draft; Writing - review & editing.

Andrea Cacciamali: Writing - review & editing.

Silvia Dotti: Writing - review & editing.

Riccardo Villa: Conceptualisation; Project administration; Supervision; Writing - review & editing.

## Data Availability Statement

All relevant data are within the manuscript.

## Ethical Approval Statement

This study did not require approval from the local ethics committee

## References

[1] W. M. S. Russell, R. L. Burch. The Principles of Humane Experimental Technique. London: Methuen & Co; 1959.

[2] Mähler M, Berar M, Feinstein R, et al. FELASA recommendations for the health monitoring of mouse, rat, hamster, guinea pig and rabbit colonies in breeding and experimental units. Lab Anim. 2014;48(3):178–192.

[3] Choi TY, Choi TI, Lee YR, et al. Zebrafish as an animal model for biomedical research. Exp. Mol. Med. Springer Nature; 2021. p. 310–317.

[4] Howe K, Clark MD, Torroja CF, et al. The zebrafish reference genome sequence and its relationship to the human genome. Nature. 2013;496(7446):498–503.

[5] Postlethwait JH, Woods IG, Ngo-Hazelett P, et al. Zebrafish comparative genomics and the origins of vertebrate chromosomes. Genome Res. 2000;10(12):1890–1902.

[6] Collymore C, Crim MJ, Lieggi C. Recommendations for Health Monitoring and Reporting for Zebrafish Research Facilities. Zebrafish. 2016;13:S138–S148.

[7] Kent ML, Harper C, Wolf JC. Documented and potential research impacts of subclinical diseases in zebrafish. ILAR J. 2012;53(2):126–134.

[8] Mocho JP, Collymore C, Farmer SC, et al. FELASA-AALAS Recommendations for Monitoring and Reporting of Laboratory Fish Diseases and Health Status, with an Emphasis on Zebrafish (Danio rerio). Comp Med. 2022;72(3):127–148.

[9] LaFollette MR, Clement CS, Luchins KR, et al. Do we still need a canary in the coal mine for laboratory animal facilities? A systematic review of environmental health monitoring versus soiled bedding sentinels. PLoS One. 2024;19(12 December).

[10] Crim MJ, Lawrence C, Livingston RS, et al. Comparison of Antemortem and Environmental Samples for Zebrafish Health Monitoring and Quarantine. Journal of the American Association for Laboratory Animal Science. 2017;56:412–424.

[11] Schuster CJ, Kent ML, Peterson JT, et al. Multi-State Occupancy Model Estimates Probability of Detection of an Aquatic Parasite Using Environmental DNA: Pseudoloma neurophilia in Zebrafish Aquaria. Journal of Parasitology. 2022;108(6):527–538.

[12] Schuster CJ, Murray KN, Sanders JL, et al. Application of an eDNA assay for the detection of Pseudoloma neurophilia (Microsporidia) in zebrafish (Danio rerio) facilities. Aquaculture. 2023;564.

[13] Kent ML, Sanders JL. Important parasites of zebrafish in research facilities. The Zebrafish in Biomedical Research: Biology, Husbandry, Diseases, and Research Applications. Elsevier; 2019. p. 479–494.

[14] Sanders JL, Lawrence C, Nichols DK, et al. Pleistophora hyphessobryconis (Microsporidia)infecting zebrafish danio rerio in research facilities. Dis Aquat Organ. 2010;91(1):47–56.

[15] Kent ML, Whipps CM, Matthews JL, et al. Mycobacteriosis in zebrafish (Danio rerio) research facilities. Comparative Biochemistry and Physiology - C Toxicology and Pharmacology. 2004;138(3):383–390.

[16] Kent ML, Feist SW, Harper C, et al. Recommendations for control of pathogens and infectious diseases in fish research facilities. Comparative Biochemistry and Physiology - C Toxicology and Pharmacology. 2009;149(2):240–248.

[17] Whipps CM, Lieggi C, Wagner R. Mycobacteriosis in zebrafish colonies. ILAR J. 2012;53(2):95–105.

[18] Astrofsky KM, Schrenzel MD, Bullis RA, et al. Comparative Medicine Diagnosis and Management of Atypical Mycobacterium spp. Infections in Established Laboratory Zebrafish (Brachydanio rerio) Facilities. 2000.

[19] Kent ML, Whipps CM, Matthews JL, et al. Mycobacteriosis in zebrafish (Danio rerio) research facilities. Comparative Biochemistry and Physiology - C Toxicology and Pharmacology. Elsevier Inc.; 2004. p. 383–390.

[20] Watral V, Kent ML. Pathogenesis of Mycobacterium spp. in zebrafish (Danio rerio) from research facilities. Comparative Biochemistry and Physiology - C Toxicology and Pharmacology. 2007;145(1):55–60.

[21] Miller M, Sabrautzki S, Beyerlein A, et al. Combining fish and environmental PCR for diagnostics of diseased laboratory zebrafish in recirculating systems. PLoS One. 2019;14(9).

[22] Leitgeb FAC, Smoak C, Horvath A, et al. The Use of Filters in the Sump for Monitoring the Health of Laboratory Zebrafish (Danio rerio). Journal of the American Association for Laboratory Animal Science. 2025;64(2):241–249.

[23] DIRECTIVE 2010/63/EU OF THE EUROPEAN PARLIAMENT AND OF THE COUNCIL of 22 September 2010 on the protection of animals used for scientific purposes (Text with EEA relevance).

[24] Slany M. A new cultivation-independent tool for fast and reliable detection of Mycobacterium marinum. J Fish Dis. 2014;37(4):363–369.

[25] Norris L, Lawler N, Hunkapiller A, et al. Detection of the parasitic nematode, Pseudocapillaria tomentosa, in zebrafish tissues and environmental DNA in research aquaria. J Fish Dis. 2020;43(9):1087–1095.

[26] Garcia KD, Coda KA, Smith AA, et al. The effects of water volume and bacterial concentration on the water filtration assay used in zebrafish health surveillance. Journal of the American Association for Laboratory Animal Science. 2021;60(6):655–660.

[27] Lawrence C, Mason T. Zebrafish housing systems: A review of basic operating principles and considerations for design and functionality. Ilar J. Oxford University Press; 2012. p. 179–191.

[28] Mocho JP. Three-Dimensional Screen: A Comprehensive Approach to the Health Monitoring of Zebrafish. Zebrafish. 2016;13:S132–S137.

[29] Batnes Matthew A. TCR. The ecology of environmental DNA and implications for conservation genetics. Conser genetic. 2016;17:1–17.

[30] Whipps CM, Matthews JL, Kent ML. Distribution and genetic characterization of Mycobacterium chelonae in laboratory zebrafish Danio rerio. Dis Aquat Organ. 2008;82(1):45–54.

[31] Yanong RPE, Pouder DB, Falkinham JO. Association of mycobacteria in recirculating aquaculture systems and mycobacterial disease in fish. J Aquat Anim Health. 2010;22(4):219–223.

[32] Tortoli E. Standard operating procedure for optimal identification of mycobacteria using 16S rRNA gene sequences. Stand Genomic Sci. 2010;3(2):145–152.

[33] Schrader C, Schielke A, Ellerbroek L, et al. PCR inhibitors - occurrence, properties and removal. J. Appl. Microbiol. 2012. p. 1014–1026.

[34] Wilson IG. Inhibition and Facilitation of Nucleic Acid Amplification. Appl Environ Microbiol. 1997;63(10):3741–3751.

